# Genomic Prediction Enables Same-Season Selection for Reduced Glycosidic Nitrile in Eastern U.S. Winter Barley

**DOI:** 10.64898/2026.06.03.729884

**Authors:** Alexis Perry, Felipe Sabadin, Wynse Brooks, Gina Brown-Guedira, Hannah Uhlmann, Harmonie Bettenhausen, Nicholas Santantonio

## Abstract

Glycosidic nitriles (GN) in barley are precursors to carcinogens formed during distillation, making GN reduction a critical breeding objective for malting and distilling industries. Measurement of GN is time-consuming. Grain must first be malted before GN can be quantified, and generally cannot be completed before selections must be made in a winter barley breeding program. Here, feasibility of same-season genomic selection against GN content was evaluated in elite Virginia Tech winter barley germplasm. In 2023, all 176 elite breeding lines screened for presence of GN were shown to be GN producers. A subset of 95 lines was then quantitatively measured for GN concentration to determine the genetic variability for the trait. Efficacy of genomic selection for GN was first assessed using a divergent selection approach on the remaining 81 predicted lines. The highest 16 and lowest 16 of the predicted lines were chosen for GN quantification. A significant phenotypic difference was found between the predicted high and low group means (0.8 ppm; *P* = 0.003). An additional 120 lines were quantified the following year to determine repeatability. GN exhibited moderate narrow-sense heritability (*h*^2^ = 0.42) and a high genetic correlation (*r* = 0.79) across years. Moderate predictive ability as was observed in cross-validation (range 0.38 − 0.61), and forward prediction using 2023 to predict 2024 (*r* = 0.39). A genome-wide scan did not identify any major-effect loci, suggesting GN content is polygenic, thus enabling same-season genomic selection to reduce GN content in this germplasm.

## 1 Introduction

As climate conditions and market demands evolve, the traits required for competitive barley varieties continue to change. Varieties that performed well 20 years ago may no longer be ideal due to the emergence of new diseases, environmental pressures, or evolving end-use quality standards (Bringhurst, 2015; Morrissy et al., 2024b). These challenges are especially relevant for barley breeding programs aiming to meet the specific needs of the malting and distilling industries.

One trait of increasing importance is the content of a family of chemicals called glycosidic nitriles (GN), which can result in the formation of ethyl carbamate (EC), a known carcinogen, during distillation. The most prevalent GN compound in barley is epiheterodendrin (EPH), which is converted during the malting and fermentation processes into EC during distillation (Bringhurst, 2015; Turner et al., 2022). GN content must be tightly managed to meet food safety and quality standards (Bringhurst, 2015). Previous work classified barley lines with GN concentrations below 0.5 ppm as GN-zero, lines ranging from 0.5 to 1.5 ppm as low-GN, and lines exceeding 1.5 ppm as high-GN based on distilling suitability and risk of EC formation (Morrissy et al., 2024b). Altering the distillation process to prevent EC formation is not a practical solution due to high costs, the risk of changing spirit flavor, and the fact that it cannot fully eliminate GN. As a result, reducing GN from the barley itself through the development of low-GN varieties represents the most effective and sustainable solution.

GN is under genetic control. Its biosynthesis is governed by a cluster of genes on chromosome 1H, including members of the CYP79, CYP71, and UGT85 gene families (Knoch et al., 2016; Ehlert et al., 2019; Jørgensen et al., 2024). In the European cultivar Emir, a large deletion within this gene cluster results in complete absence of GN production (Ehlert et al., 2019). Detecting this GN-null deletion through traditional phenotyping methods can be time-consuming and costly, prompting the development of SNP markers linked to the deletion to enable more efficient identification of GN-zero lines (Swanston et al., 1999; Ehlert et al., 2019).

Introgressing GN-null alleles into elite germplasm provides a long-term solution for eliminating GN production, but selecting for reducing GN levels within elite breeding material is crucial to prevent release of varieties with high GN production. The breeding cycle for winter barley can require ten to twelve years from the initial cross to variety release, and backcrossing into adapted germplasm only prolongs this process. Therefore, there is a need for tools that can identify and advance low-GN lines while GN-zero populations are under development.

Understanding the genetic basis of GN is essential for integrating GN as a consistent breeding target, but phenotyping for GN remains a major bottleneck. In winter grains, where planting occurs only two to three months after harvest, breeding decisions must generally be made before GN phenotypes can be measured. GN phenotyping of large numbers of samples is time-consuming due to the need to malt small samples of grain (250 g - 350 g) for each line before GN can be quantified. Depending on the laboratory and analytical procedure, GN quantification can be very costly, ranging from $100 to $225 per sample (Ehlert et al., 2019). This limits the ability to directly select for reduced GN content in early generations where the number of candidate lines is large.

Genomic selection, therefore, may be a useful strategy to perform same-season selection on early generation materials, provided they are genotyped with a sufficiently dense marker panel (a common practice in modern breeding programs), and sufficient GN phenotyping has already been performed on related materials. Genomic selection uses genome-wide marker data to predict genomic estimated breeding values (gEBVs) for individuals without requiring phenotypic data on every line (Meuwissen et al., 2001; Bernardo, 1994;). A genomic prediction model is “trained” by using a population that has been both phenotyped and genotyped, estimating marker effects (typically in a mixed model framework), and using those marker effect estimates to predict the genetic merit of related, but unobserved lines that have also been genotyped. In a winter barley breeding program aimed at reducing GN, new early generation materials (e.g. first year of yield testing) can be predicted as soon as genotypic information and phenotypes from the previous year are available. Then lines with low predicted GN may be advanced into later stages of testing and validated after trials have been planted, while lines with high predicted GN may be dropped. Genomic predictive ability depends on several factors, including the heritability of the trait, the extent of linkage disequilibrium between markers and causal loci, and the size and relatedness of the training population (Sallam et al., 2015; Ould Estaghvirou et al., 2013).

The objectives of this study were to (1) characterize genetic variation of GN levels within Virginia Tech’s (VT) elite barley germplasm, (2) evaluate the potential for using genomic prediction to identify lines with reduced GN content, and (3) determine whether genomic prediction could support same-season selection for reduced GN content in winter barley breeding programs. Lines were first screened using a colorimetric assay to determine the presence or absence of detectable EPH and identify whether any VT germplasm carried a GN-null allele. A subset of these lines was then selected for quantitative measurement of GN levels and development of genomic prediction models, which were in turn used in a divergent selection approach as a proof of concept. Predictive ability was evaluated using cross-validation within and across years to determine whether genomic data could reliably support same-season selection for reduced GN content in winter barley breeding programs.

## 2 Materials and Methods

### 2.1 Plant Materials and Field Experiment Design

This study utilized elite two-row breeding lines from the Virginia Tech barley breeding program, defined as lines in their second year or beyond of agronomic evaluation. In 2023, a total of 176 elite lines, along with two known GN-zero checks (178 lines total), ‘Top Shelf’ (Morrissy et al., 2024a) and ‘GN0-Vivar’, from Oregon State University, were evaluated. In 2024, 39 lines were advanced from 2023 trials and 81 new lines were added, resulting in a total of 120 lines evaluated during the second year of this study. No selection on GN was performed for advancement of the 39 lines.

Lines were grown in a series of replicated field trials representing different stages of the Virginia Tech barley breeding pipeline. These included the Hulled Preliminary nursery, consisting of hulled barley lines in their second year of agronomic evaluation; the Malt Quality nursery, consisting of lines with moderate agronomic performance but good malt quality characteristics; and the Eastern Malt nursery, an advanced regional trial containing the highest-performing breeding lines.

The Hulled Preliminary and Malt Quality trials were evaluated in Blacksburg, VA and Warsaw, VA using two replicates per location. The Eastern Malt trial was also evaluated in Blacksburg, VA, and Warsaw, VA, with three replicates per location, as well as other locations in the Eastern U.S. (data not reported). Although lines were evaluated across multiple locations in teh Eastern Malt nursery, grain samples used for glycosidic nitrile (GN) phenotyping were collected only from Blacksburg and Warsaw.

### 2.2 Genotyping

All lines included in this study were genotyped using genotyping-by-sequencing (GBS) at the USDA-ARS Wheat Genotyping Lab in Raleigh, North Carolina. Genomic DNA was digested using a two-enzyme GBS protocol with the restriction enzymes *Pst1* and *Msp1* (Elshire et al., 2011). GBS libraries were sequenced using Illumina sequencing technology, and variant calling was conducted using the TASSEL pipeline aligned to the Morex v3 barley reference genome via the BWA alignment algorithm (Bradbury et al., 2007; Mascher et al., 2021; Li and Durbin, 2010). Initial raw variant data included 810,451 single-nucleotide polymorphisms (SNPs) across 2,186 individuals.

Variants were then filtered using BCFtools to exclude markers with more than 30% missing data, more than 10% heterozygosity, or a minor allele frequency less than 0.01 (Danecek et al., 2021). Missing genotype calls were imputed using Beagle (version 5.2) (Browning et al., 2021). After filtering, imputation and a filter for markers in very high LD (*r >* 0.99), the final genotype matrix included 22,587 high-quality SNPs. This curated GBS data was subset of the individuals included in the study, and used for both genomic prediction.

### 2.3 Glycosidic Nitrile Phenotyping

Glycosidic nitrile phenotyping consisted of a two-stage process involving qualitative screening for epiheterodendrin (EPH) followed by quantitative GN determination. EPH is the principal GN present in barley and serves as an indicator of GN production potential. There is a less expensive colorimetric test for the presence of EPH that can identify lines homozygous for GN-null alleles, but it cannot quantify the amount of GN in lines that do express it. Briefly, malt extracts are reacted with assay reagents that produce a colorimetric response in the presence of EPH, allowing samples to be classified as positive or negative for detectable EPH (Xue et al. 2023). To determine if any GN-zero lines containing the GN-null allele existed in the breeding program, it was decided to screen all elite lines with this colorimetric EPH assay first. Grain samples from 178 elite barley lines sourced from Blacksburg, VA, in 2023, including two known GN-zero checks from Oregon State University, were screened for EPH at the Hartwick College Center for Craft Food and Beverage in Oneonta, New York following the methods described by Xue et al. (2023).

Other than the two known GN-zero checks, no GN-zero individuals were identified within the Virginia Tech barley germplasm during initial screening. Therefore, a subset of 95 lines, including the two GN-zero checks, were further quantified for GN levels. These lines were chosen to represent the diversity of breeding material evaluated across the trials while capturing the observed range of EPH assay responses present within the population. Quantitative GN concentrations were determined according to EBC Analytica 4.21 Glycosidic Nitrile (European Brewery Convention, 2006) and reported as parts per million (ppm). Because samples had to be malted before GN was quantified, other malt quality characteristics were also measured, including malt extract, *β*-glucan (BG), soluble protein (SP), free amino-nitrogen (FAN), diastatic power (DP) and *α*-amylase (AA). After genomic predictions were made, an additional 32 samples were then quantified as part of a divergent genomic selection approach (see ‘Divergent Genomic Selection’ below).

In 2024, 120 lines were subjected to quantitative GN analysis to evaluate repeatability across years. Samples were initially intended to be analyzed at Hartwick College to maintain consistency with the 2023 dataset. However, staffing turnover and equipment malfunction at Hartwick College necessitated transfer of the analyses to the Montana State University Malt Quality Lab in Bozeman, Montana. Additionally, excessive rainfall and subsequent lodging reduced grain quality in Blacksburg during 2024. As a result, grain samples used for GN quantification were collected from Warsaw, VA, where field conditions were more favorable. Both years included the GN-zero checks as controls. As such, repeatability was assessed not only across years, but also across locations and analytical labs.

Phenotypic reproducibility across years was evaluated using Pearson correlations (*r*) between GN measurements of lines evaluated in both the 2023 and 2024 datasets. The genetic correlation across years was evaluated by a Pearson correlation of gEBVs for lines measured in both years. Phenotypic and genetic correlations among GN and malt quality traits were estimated using Pearson correlation coefficients.

### 2.4 Genomic Prediction Methodology

Genomic prediction was conducted using genomic best linear unbiased prediction methodology (gBLUP; equivalent to a ridge-regression on markers), and fit using the rrBLUP package in R (Endelman, 2011).

The following mixed linear model was used:

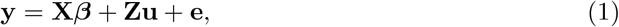

where **y** is the vector of phenotypic observations for GN content, **X** is the design matrix for fixed effects, ***β*** is the vector of fixed effects, **Z** is the incidence matrix relating observations to line identities, **u** is the vector of random additive genetic effects with covariance, **K**, and **e** represents residual error (Yu et al., 2006). For analyses conducted within each year, the fixed effect, ***β***, was simply an intercept, while a year predictor was added to the across year analyses. The kinship matrix (***K***) was calculated using centered and scaled GBS marker data, according to the method described by VanRaden (2008), to represent the additive genetic covariance among lines.

### 2.5 Divergent Genomic Selection

As a proof-of-concept, a divergent genomic selection approach was implemented to determine if significant gains evaluation of same-season selection utilizing a was implemented. The GN values for the 95 phenotyped lines from 2023 were used as the training population to generate genomic predictions for the 83 unphenotyped lines from that year. Predicted GN values were ranked from lowest to highest. The sixteen lines with the lowest predicted GN values and the sixteen lines with the highest predicted GN values were selected and for quantitative GN analysis. Selected lines were randomized prior to laboratory analysis to minimize potential batch effects during GN quantification. The resulting 32 validation lines, together with the original 95 and the 120 lines quantified in 2024, were included in subsequent genomic predictive ability analyses.

### 2.6 Predictive Ability and Heritability

Predictive ability was evaluated using multiple validation strategies: (1) ten-fold cross-validation within the 2023 phenotyped dataset, (2) ten-fold cross-validation within the 2024 phenotyped dataset, (3) ten-fold cross-validation across the combined 2023 and 2024 dataset, and (4) forward prediction, where models trained on 2023 phenotyped lines were used to predict GN values in 2024 lines. Predictive ability was quantified as the Pearson correlation coefficient (*r*) between observed GN values and predicted gEBVs.

Narrow-sense heritability (***h***^**2**^) was also estimated for GN from variance component es-timates from the fitted model as:

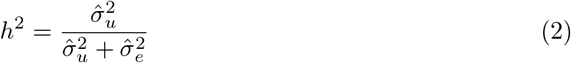

Where 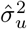 represents additive genetic variance estimate and 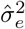 represents residual variance estimate. These estimates provide insight into the proportion of phenotypic variance attributable to additive genetic effects and help evaluate the expected response to selection. Together, predictive ability and heritability estimates inform the feasibility of integrating genomic prediction into breeding strategies targeting GN reduction (Fisher, 1918; Schmidt et al., 2019; Puglisi et al., 2021).

## 3 Genome-wide scan

A genome-wide scan for marker trait associations was conducted using rrBLUP in R (Endelman, 2011) on data collected in both years to determine if loci with large effects on GN content existed within the Virginia Tech breeding germplasm. Although the primary objective of this study was genomic prediction and same-season selection, this scan was conducted to inform if a marker-assisted selection approach would be more effective than a genomic selection approach. A Bonferroni correction was used as a genome-wide significance threshold.

## 4 Results and Discussion

### 4.1 Phenotypic Variation in GN Levels

The EPH assay screening confirmed that the only GN-zero individuals present were the known GN-zero checks from Oregon State University. GN levels ranged from 0.3 to 5.3 ppm with a mean of 2.6 ppm (Figure S1a). In 2024, GN levels for 120 elite VT lines ranged from 0.2 to 4.6 ppm, with a mean of 2.0 ppm (Figure S1b).

The absence of GN-zero individuals within the VT breeding germplasm indicated that the EPH-null allele was not present among the elite breeding lines evaluated in this study. However, substantial variation in quantitative GN levels was observed within the population, suggesting that sufficient phenotypic variation exists to support selection for reduced GN content.

### 4.2 Bidirectional Genomic Selection Validation

Genomic prediction models trained using 95 phenotyped lines were used to estimate gEBVs for 83 unphenotyped lines. Predicted GN values were ranked from lowest to highest and used to identify 32 candidate lines for divergent genomic selection (Figure 1a)

**Figure 1:**
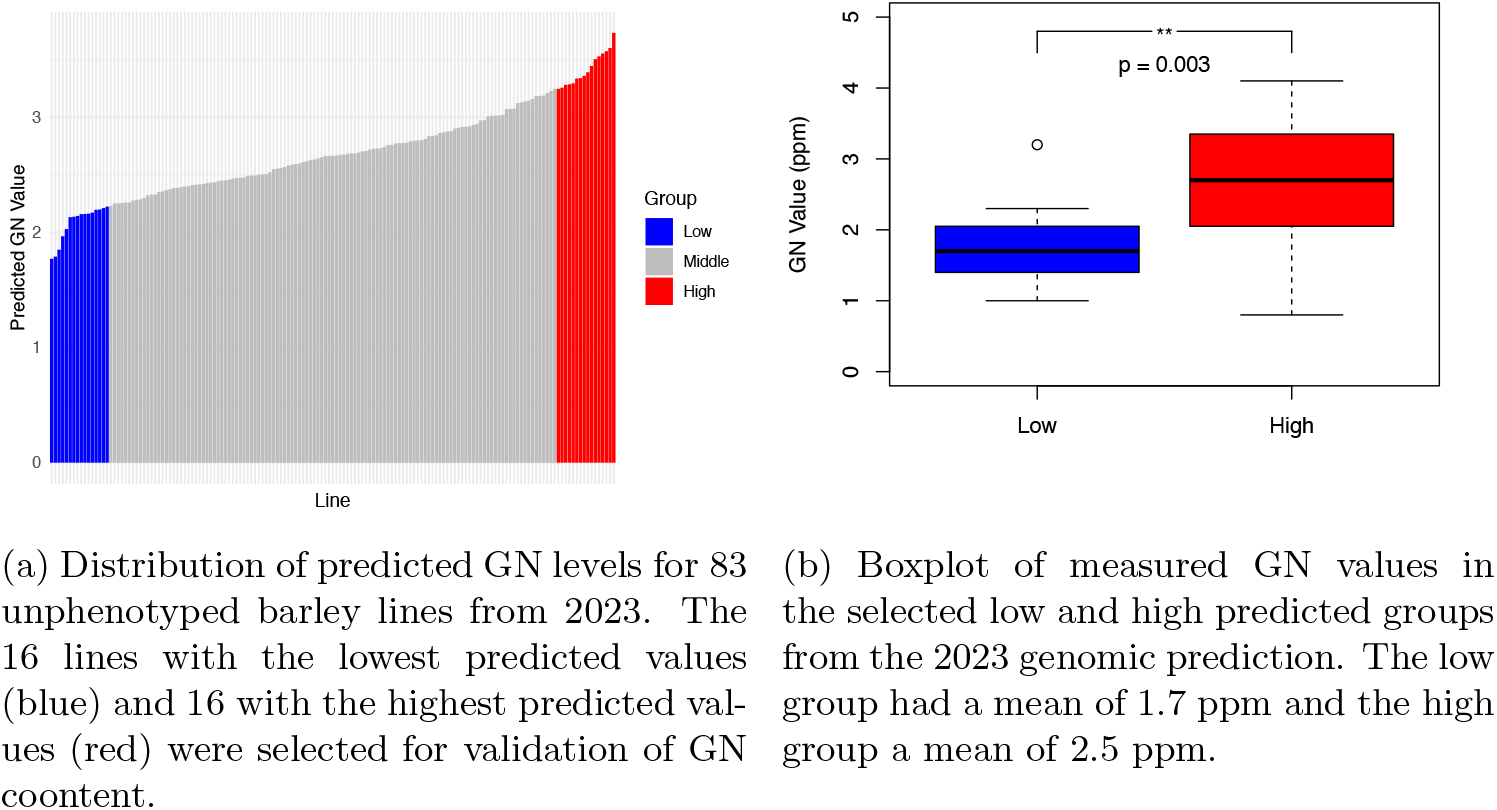
Genomic predictions and validation of divergent selection of GN levels in Virginia Tech barley lines from 2023.

Measured GN values differed significantly between the predicted high- and low-GN groups as determined by a t-test (*P* = 0.003). The low-predicted group had an observed mean GN value of 1.7 ppm, while the high-predicted group averaged 2.5 ppm (Figure 1b).

Scatterplots comparing predicted and measured GN values showed moderate agreement between predicted and observed GN levels, although overlap between groups remained (Figure S2). This proof of concept demonstrated that despite relatively low heritability observed in 2023 (h = 0.25; see Genomic Prediction Performance section below), genomic selection was effective.

### 4.3 Environmental Effects on GN Expression

While the proof of concept was successful, it was unclear if large genotype by environmental effects for GN content across years or locations would negate the genomic selection appraoch. Thus repeatability across years was assessed by evaluating lines the following year in a separate location. A phenotypic correlation of *r* = 0.73 was observed between measured GN values of 39 lines evaluated in both years (Figure 2a). The corresponding genotypic correlation was slightly higher at *r* = 0.79 (Figure 2b).

**Figure 2:**
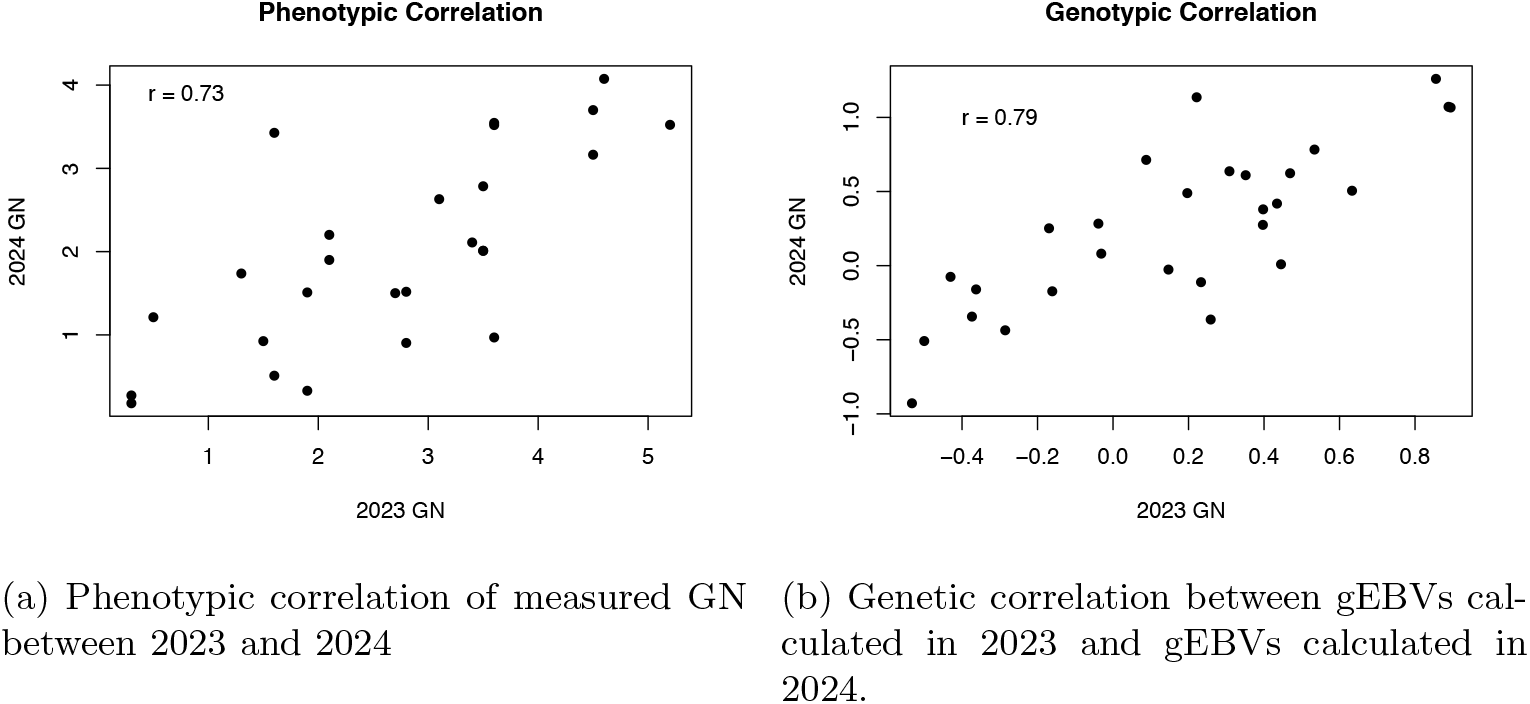
Year-to year phenotypic and genetic correlation of GN for 39 lines evaluated in both 2023 and 2024.

GN showed significant positive phenotypic correlations with soluble protein (*r* = 0.25, *P* = 0.0002) and free amino nitrogen (*r* = 0.24, *P* = 0.0002), a significant negative correlation with *β*-glucan (*r* = −0.21, *P* = 0.001), and no significant relationship with diastatic power (*r* = 0.11, *P* = 0.1). Genotypic correlations were mostly stronger, for example, with GN and *β*-glucan showing a moderately strong negative correlation (*r* = −0.47, *P* = 0.0001) (Figure 3). With the exception of *β*-glucan, none of these correlations are strong enough to exhibit concern that selection for low GN would result in indirect selection that would reduce malting quality.

**Figure 3:**
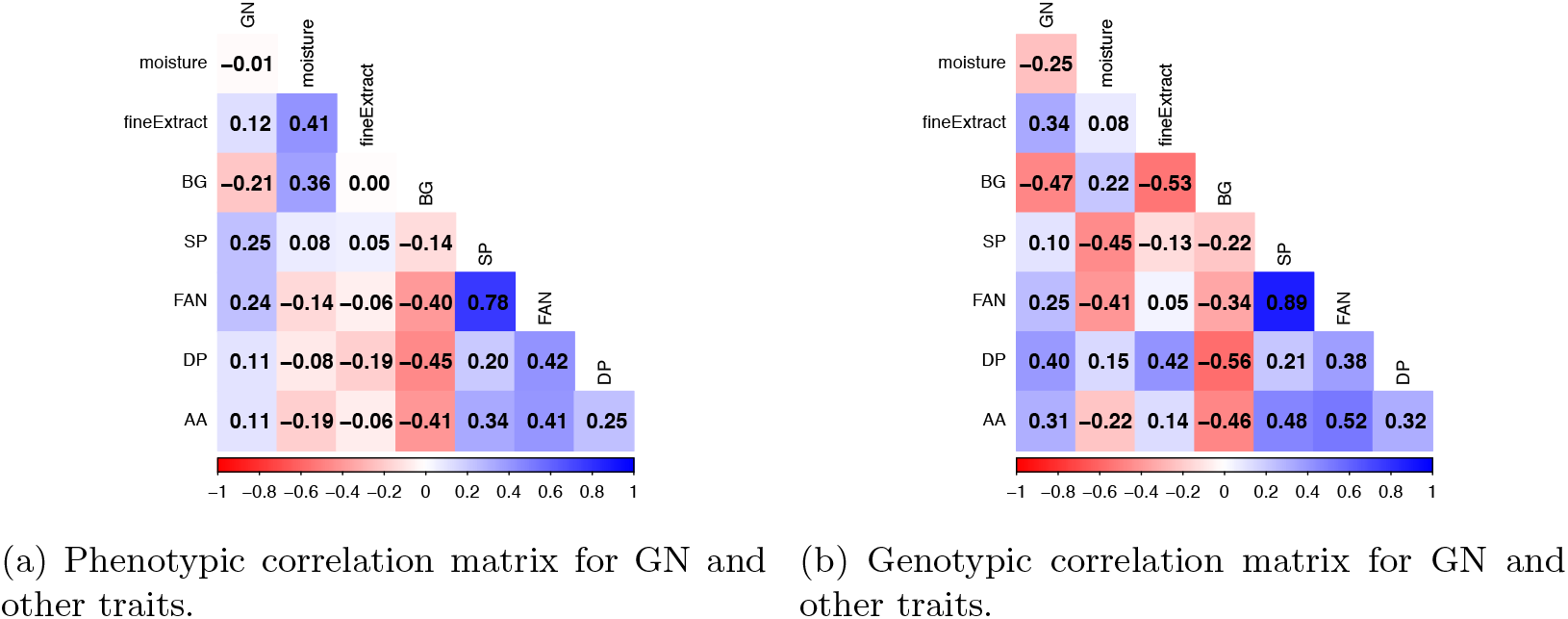
Phenotypic and genetic correlations for GN and other malting quality traits. GN: glycosidic nitrile, BG: *β*-glucan, SP: soluble protein, FAN: free amino nitrogen, DP: diastatic power, AA: *α*-amylase.

GN was negatively associated with *β*-glucan in both years, although the strength of the relationship differed between datasets (Figure S3). One important challenge in GN phenotyping is differentiating genetically low-GN lines from lines with artificially low GN values caused by under-modification during malting. Elevated *β*-glucan can indicate poor modification and may result in GN measurements that do not reflect true genetic differences. Samples from 2023 appeared to have some lines that were under-modified, evidenced by a stronger relationship between *β*-glucan and GN, due to either an environmental effect or a difference in lab malting procedure. Future phenotyping efforts should integrate malt quality indicators, such as *β*-glucan and *α*-amylase, alongside GN quantification to improve selection accuracy.

GN levels were quantified at Hartwick College in 2023 and Montana State University in 2024 due to unforeseen logistical issues. Differences in equipment, reagents, or handling may have introduced variability between years. However, the extent to which laboratory differences contributed to reduced correlations across years cannot be separated from environment or location effects. The relatively strong phenotypic and genotypic correlations observed across years nevertheless suggest that GN expression remains under substantial genetic control despite environment and procedural variation.

### 4.4 Genomic Prediction Performance

Predictive ability ranged from 0.38 to 0.61 within years, 0.52 across the combined dataset, and 0.39 for forward prediction (Table 1). Despite moderately low heritability estimates within year (*h*^2^ = 0.25 in 2023 and *h*^2^ = 0.35 in 2024), genomic prediction demonstrated useful predictive ability for GN content. Predictive ability increased when the larger combined dataset was used, suggesting that additional phenotypic observations improved model performance, which is not surprising given the relatively small size of the dataset (247 lines measured for GN). The ability of the model to maintain predictive ability across years despite differences in growing environments (year and location) and analytical laboratories suggests that GN expression is sufficiently stable to support same-season selection in winter barley breeding programs.

**Table 1:**
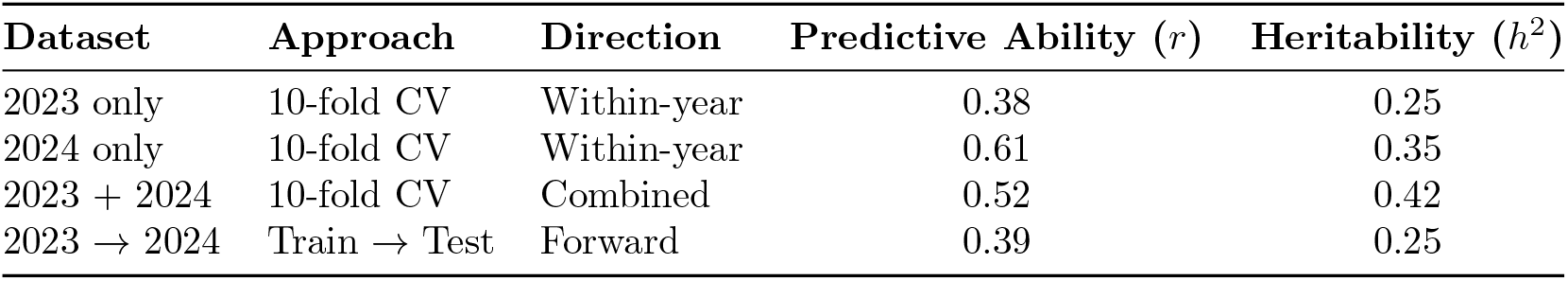
Predictive ability (*r*) and narrow-sense heritability (*h*^2^) for GN content across genomic prediction scenarios. Accuracy values are shown separately for methods using five-fold cross-validation (CV) or forward prediction (Train → Test) across datasets.

Predicted GN values generated using the 2023 training population were moderately correlated with measured GN values in the independent 2024 dataset (Figure 4). The 39 lines evaluated in both years generally followed the same overall trend as newly evaluated 2024 lines, indicating that genomic prediction remained informative across years and environments.

**Figure 4:**
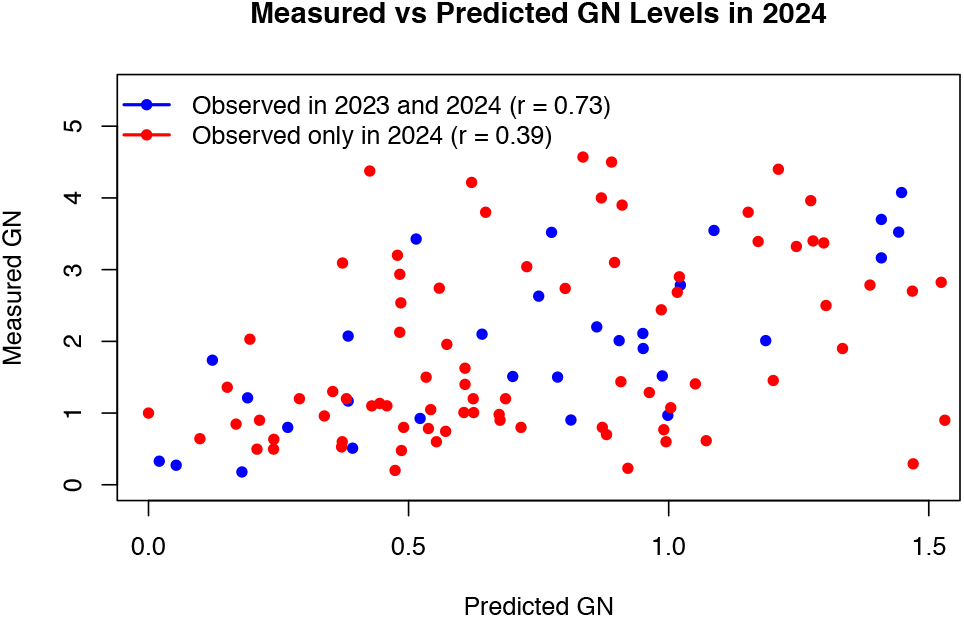
Relationship between gEBVs for GN generated from the 2023 training population and measured GN values observed in 2024. Blue points represent lines evaluated in both 2023 and 2024, while red points represent lines evaluated only in 2024.

### 4.5 Genome-wide scan

Genome-wide association analysis conducted on GN measured in both years did not identify any SNPs surpassing the genome-wide significance threshold (Figure S4). The absence of significant marker associations likely reflects the limited number of GN-zero lines present in the analysis (two GN-zero checks) and the relatively small number of phenotyped individuals (247). Although no markers exceeded the Bonferroni-correct significant threshold, there were a number of markers on chromosome 1H with low p-values near the previously reported EPH region. This suggests allelic variation in and around the EPH locus may still be contributing to genetic variation for GN, despite the absence of the GN-null allele in the population. Several other locations in the genome showed suggestive peaks that did not pass the significance threshold.

## 5 Conclusion

The study evaluated glycosidic nitrile (GN) levels in elite Virginia Tech barley breeding lines and assessed whether GN can be predicted using genomic data to support breeding strategies for GN reduction. Despite a modest narrow-sense heritability estimate (*h*^2^ =0.25) genomic prediction effectively differentiated high- and low-GN lines following genomic divergent selection. Strong correlation in GN expression across years, environments, and analytical laboratories further supported GN as a viable breeding target. Altogether, these findings demonstrate that genomic prediction can support same-season selection for GN in winter barley breeding programs. This work provides a practical validation that genomic selection of grain quality traits is an effective approach, especially when they cannot be easily measured for same-season selection. This process is now being used at Virginia Tech to select for low-GN barley varieties suited to distilling markets while GN-null alleles are advanced into elite Virginia Tech breeding materials.

## Supplementary Materials

**Figure S1:**
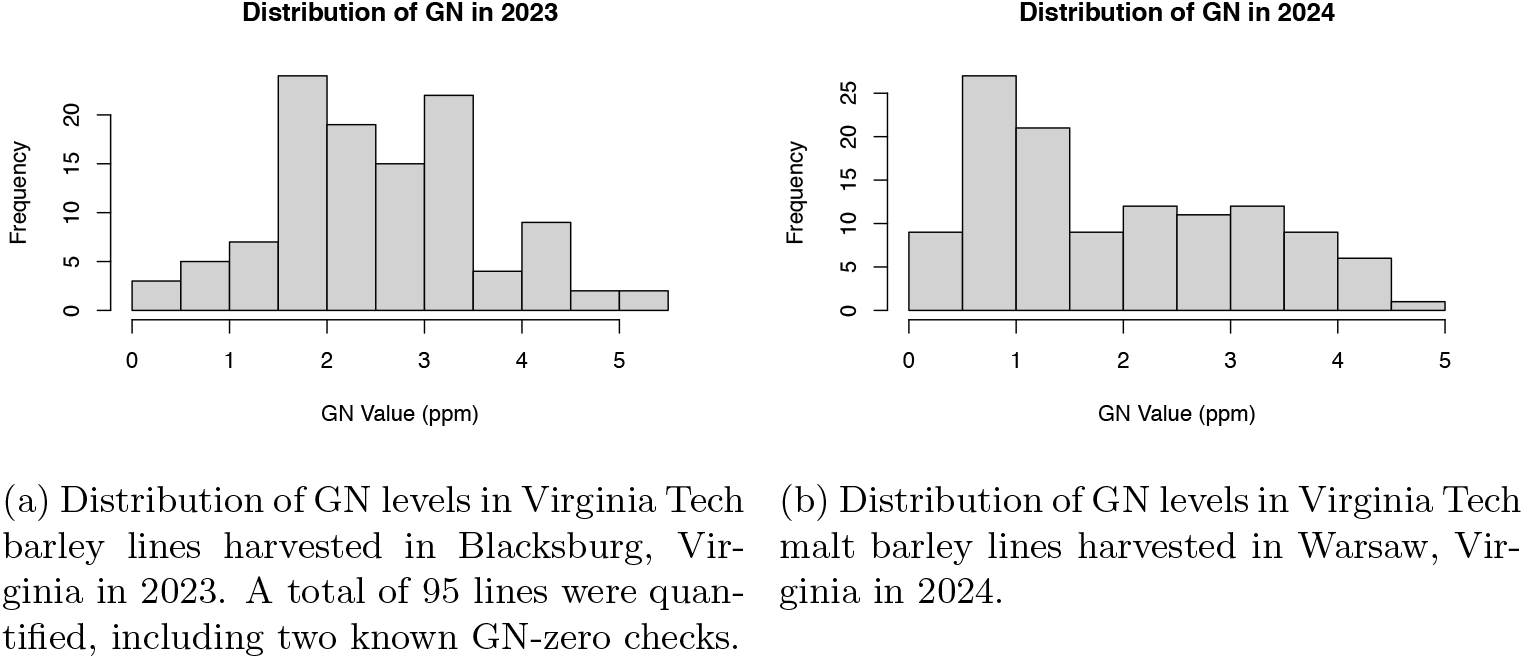
Distribution of GN levels in Virginia Tech barley lines evaluated in 2023 and 2024.

**Figure S2:**
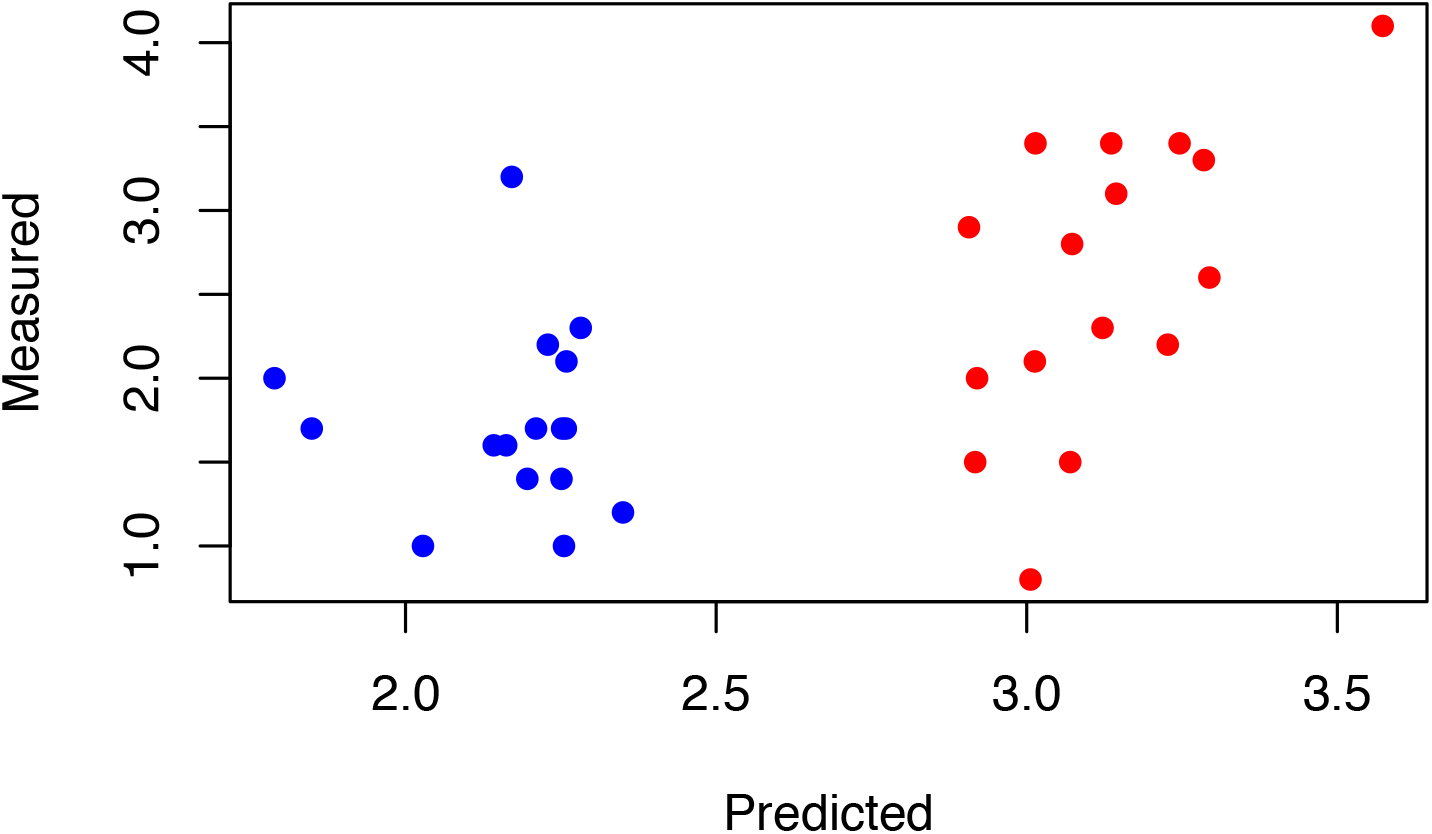
Scatterplots comparing predicted and measured GN values (ppm) for validation lines selected in 2023 with the low lines highlighted in blue and the high in red.

**Figure S3:**
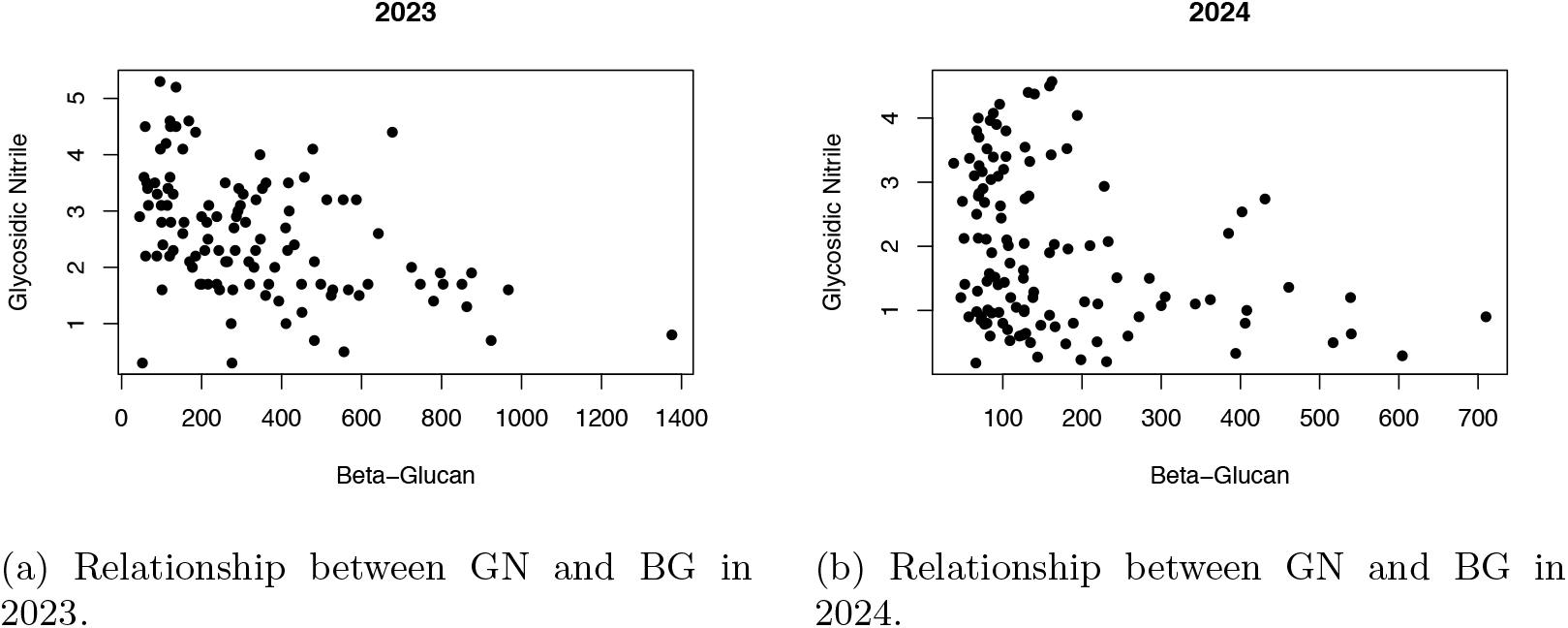
Scatterplots of GN versus *β*-glucan (BG) content in 2023 and 2024, highlighting year-to-year trends in modification levels.

**Figure S4:**
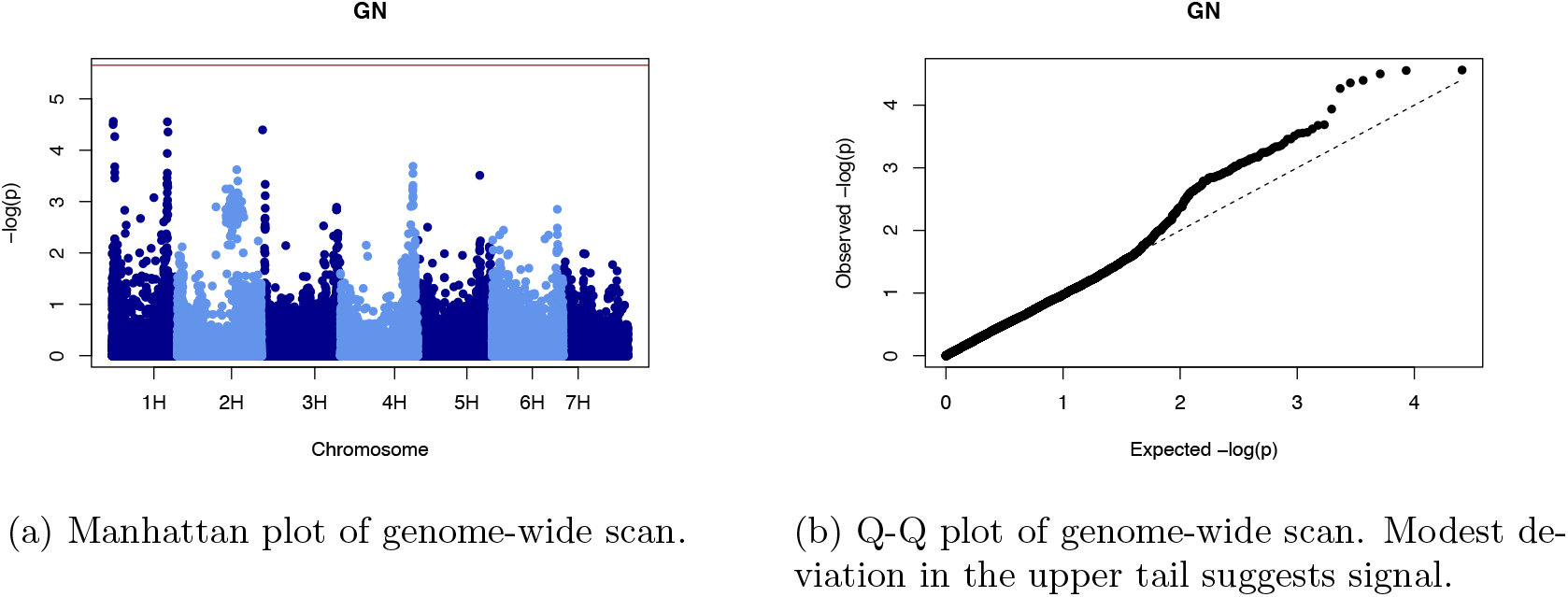
Genome-wide scan results.

